# A Retina-Inspired Computational Model for Stimulation Efficacy Characterization and Implementation Optimization of Implantable Optogenetic Epi-Retinal Neuro-Stimulators

**DOI:** 10.1101/2023.03.28.534592

**Authors:** Tayebeh Yousefi, Hossein Kassiri

## Abstract

In this paper, a biologically-informed computational framework is developed to model the efficacy and to optimize the implementation of an implantable epi-retinal prosthesis that performs optogenetic stimulation through a *μ*LED array. The developed model is capable of translating visual stimulus inputs into corresponding signals evoked in the transfected retinal cells through optogenetic stimulation, calculating the subsequent neuronal activities of the following retinal layers, and estimating the resulted brain’s visual perception. As such, it can model and quantitatively analyze the impact of optical stimulation parameters (intensity, frequency, directivity, wavelength, etc.) and the *μ*LED array’s physical specifications (array size, density, pitch, implantation location, etc.) on the efficacy of the stimulation.

Using this model, we compared optical and electrical stimulations in terms of the structural similarity between their induced visual perception in the brain and the visual stimulus input. We showed that thanks to the cell-type specificity of optogenetic stimulation, it can induce more relevant visual perception qualities than electrical stimulation. We also showed that its resulted visual perception substantially improves with scaling the stimulator’s array size. The model was also used to qualitatively and quantitatively analyze the impact of parameters such as implantation location, light intensity, single- and dual-wavelength stimulation, and illumination divergence angle on the quality of the optical-stimulation-induced visual perception. In each case, the simulation results were followed by our interpretation from a biological point of view. More importantly, in each case, we discussed how the results could be used for optimizing different parameters of an implantable optogenetic stimulator to achieve maximum efficacy and energy efficiency. Keywords: Retinal prosthesis, optogenetics, visual perception, optical stimulation, *μ*LED array, computational model, spatial resolution, pathway-specific stimulation.

## I. Introduction

As the leading cause of vision loss, age-related macular degeneration (AMD) affects 196 million people worldwide— a number that is projected to double by 2050 [1], [2]. AMD patients gradually lose their vision due to the degeneration of photo-receptor layer cells in their retina. Regardless of the underlying reason being environmental, pathologic damage, or genetic mutation, currently this progressive neurological disorder has no established cure, and various pharmacological therapies can only slow down its progression [3]. Over the past few decades, a variety of multi-disciplinary approaches, including electronic retinal prostheses, optogenetics, chemical neural interfaces, gene therapy, and cell transplantation have been used in an attempt to restore visual function through bypassing the damaged photo-receptor layer [4]–[7]. Most of these methods rely on the fact that despite the photo-receptors deterioration and reorganization, the inner retinal neurons largely retain their capacity for signal transmission [3].

As shown in Fig. 1 the mammalian retina is located at the back of the eyeball and consists of rod and cone photo-receptors, horizontal cells, bipolar cells, amacrine cells, and retinal ganglion cells (RGCs). It is the axons of RGCs that are bundled together to form the optic nerve, which is responsible for transmitting visual information from the retina to the brain. As illustrated by the “visual stimulus” direction in the picture, the light passes through layers of neurons before activating the rod and cone photo-receptors. In a healthy retina, upon receiving an incident light and proportional to its power, photo-receptors generate photo-currents that trigger the downstream retinal network indicated by the “visual information stream” direction in Fig. 1. For a degenerated retina, the loss of functionality in photo-receptors prevents the first stage of this chain from generating visually-evoked signals, which results in loss of vision despite the fact that the downstream network of neurons (i.e., horizontal, bipolar, amacrine, and RGCs) is healthy and functional.

**Fig. 1:**
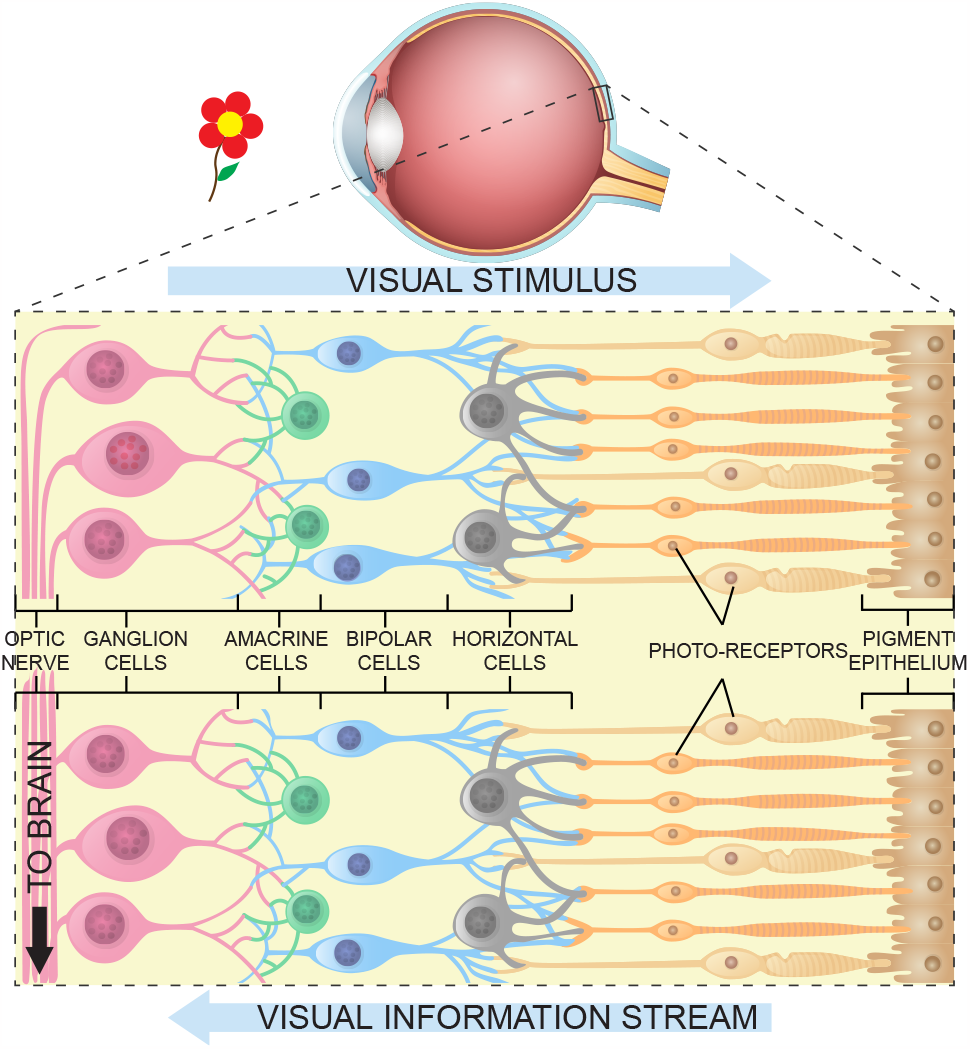
The structure of different layers in a mammalian retina, illustrating the visual input light path and the visual data processing network.

The downstream network acts as a series of parallel information processing paths, where different features of the stimulus image are extracted by various types of cells in each layer. The RGCs are the last layer of the processing path and each RGC cell-type is activated by a specific feature extracted from the visual stimulus. In recent years, implantable electronic retinal prostheses have shown potential to offer a promising treatment option for retinal degeneration by manipulating its response to light through artificial stimulation of the above-mentioned neuronal layers that carry visual information to the brain [8]–[13]. In an electronic retinal prosthesis, the visual stimulus is captured using an image sensor (typically, a camera placed on a goggle or an eye glass) and is sent to an external processing unit to generate corresponding stimulation patterns for artificial electronic stimulation of the inner retina layers (i.e., bipolar cells or RGCs). As shown, the external unit wirelessly communicates these commands to an implantable chip, where it is decoded to generate the control and timing signals required for conducting the stimulation. Based on the placement of the implant with respect to the retina, the electronic retinal prostheses are divided into two main categories of epiretinal and sub-retinal.

The epi-retinal prosthesis is attached to the innermost layer of retina (i.e., the RGCs), which is the most common placement option, mainly thanks to its simpler surgical implantation [8]–[11]. However, since they are in direct contact with RGCs (i.e., the last layer in the retinal downstream network of neurons), all other healthy layers (i.e., bipolar, amacrine, etc.) are also bypassed. As mentioned previously, the job of these bypassed layers is to extract and accentuate features (i.e., visual information) from the photo-receptors’ output and to pass along visual information and generate corresponding spiking patterns at RGC layer to relay the message to the brain. In the absence of these layers, this job needs to be done by the electronic circuits, using image processing techniques or artificial neural networks, implemented on-chip to mimic the functions of the bypassed downstream neuronal network [14]–[16]. Due to the complexity of the bypassed network, the cross-patient variations, and the limited computational resources, the implementation of the image processing unit is not efficient nor practical. In fact, the full extent of image processing that is taken place in the retina downstream network has not been fully recognized and there are more than 20 types of RGCs, each assumed to convey a different aspect of visual information.

Motivated by this, sub-retinal stimulators are placed on the other end of retinal structure, where the degenerated photo-receptor layer is located. This type of prostheses can target the middle layer of retina (i.e., bipolar layer) and rely on the downstream retina network to perform the signal processing in the natural way. As such, stimulation pulses only need to mimic the photo-receptors’ outputs, which are proportional to the incident light sensed by the image sensor— a significantly more feasible task to do compared to epi-retinal stimulators’. However, the implantation surgery for sub-retinal implants are more challenging and it can cause severe inflammation and implant rejection. Additionally, the limited available space for the implant and being away form vitreous gel which has temperature regulatory feature, result in tight restrictions on implant’s size and heat dissipation, respectively[12], [13], [17].

Besides the above-described challenges, a fundamental barrier that severely limits electrical (voltage or current mode) stimulators’ efficacy in restoring vision is their indiscriminate activation of all types of retina cells that are in the proximity of stimulation electrodes. As will be described in detail in section II, this could result in contradicting messages being communicated to the brain through different neural pathways, resulting in a poor or noise-like image reconstruction [18], [19]. In an effort to address this, and to improve visual perception’s quality and consistency, neuro-chemical retinal prostheses are proposed (e.g., [20]) that employ certain chemical neurotransmitters to discriminately target different types of retinal cells to achieve a target stimulation pattern. However, delivery of these chemical substances in a spatially- and temporally-controlled fashion poses significant challenges that do not bode well with stimulation channel-count scaling, thus preventing the development of a multi-channel implantable device capable of performing this task to date.

In this work, through development of a customized computational model, we will investigate the feasibility of employing an implantable wireless optical neurostimulator (e.g., our recently-reported device in [21]) as a treatment option for patients with retinal degeneration, and compare its performance and potential to an electrical stimulator from various aspects. This is mainly motivated by the optogenetics’ inherent cell-type specificity that enables pathway-specific activation of neurons, allowing for a visual perception quality that is practically unachievable with electrical stimulation. Optogenetic stimulation requires genetic modification of the degenerated retinal cells using microbial opsins to restore their light sensitivity. While this has been done in the past, it has been shown (e.g., in [7]) that the opsins’ sensitivity to ambient light is far less than a healthy eye’s photo-receptors. As such, we propose to use an implantable optical stimulator to activate genetically-modified opsin-transfected cells of a retina layer (i.e., bipolar cells or RGCs). Thanks to the light transparency of the retina’s neuronal layers, the optical stimulator could be placed on the RGC side (i.e., easier surgical implantation), while targeting one of the outer layers in retina (e.g., bipolar), thus can take advantage of the healthy downstream retina network to perform the signal processing.

The presented customized computational model is developed based on a widely-used framework for modeling human vision and is designed to be capable of (a) translating a given visual stimulus input to optical stimulation patterns, (b) estimating the brain’s visual perception for a given optical stimulation pattern, and (c) evaluating the impact of varying optical stimulation parameters (e.g., *μ*LED array size, light intensity, light wavelength, illumination divergence angle, etc.) on the inferred perception. Using this model, we have compared optogenetic and electrical stimulation performance and potential for an implantable medical device aimed at restoring vision in patients with retinal degeneration. The quantitative comparison is made in terms of the quality of the inferred visual perception, the scalability of the *μ*LED/*μ*electrode array size and density, and the corresponding achievable spatial resolution.

The model is also used to investigate the sensitivity of the visual perception to various optical stimulation parameters (e.g., *μ*LEDs’ type/wavelength, array pitch and density, etc.). Optimization of these parameters is crucial for the design of an implantable retinal prosthesis under a tight power budget, where energy consumption needs to be minimized without sacrificing the stimulation efficacy. Besides, considering that an acceptable visual perception by the brain requires scaling up the number of stimulation channels to large quantities, a suboptimal solution could result in excessive heat generation, stimulation of blind spots, visual perception saturation, or insufficient spatial resolution and/or coverage. We present this work as a primary step towards finding and optimizing key design parameters and requirements of an implantable optogenetic-based retinal prosthesis. This paper extends on an earlier report of the principle and demonstration in [24], and offers a more detailed description and analysis of the presented model, as well as additional comparative discussions and simulation results on parameter optimization.

The rest of the paper is organized as follows. Section II explains the significance of cell-type-selective activation in a retinal prosthesis and highlights the advantageous of optogenetic stimulation in this regard, compared to electrical stimulation. Section III describes the development details of the presented computational model. It also includes a comparative discussion on the efficacy of the electrical and optogenetic stimulations based on the model’s simulation results. Section IV investigates the effect of different optical stimulation parameters and the *μ*LED array’s physical specifications through conducting various simulations and discusses their implications in optimizing the design of a retinal prosthesis in terms of biological efficacy, spatial resolution, and energy efficiency. Section V concludes the paper.

## II. Cell-type Specificity

### A Requirement in Retinal Prostheses

As mentioned, the downstream retinal chain (from photo-receptors to RGCs) acts as a series of parallel signal processing network with more than 20 types of RGCs whose receptive fields tile the retinal surface and convey distinct visual information about the input visual stimulus [25]. In a healthy retina, the changes in the power of incident light and its associated photocurrent trigger two morphologically distinct pathways, called ON and OFF pathways, which are activated by increments or decrements of light, respectively. Fig. 2 shows how the parasol and midget cells (two types of RGCs) tile the surface of the RGC layer. These numerically dominant cell types are further divided into ON and OFF sub-types, which respond antagonistically to the changes in light intensity. Under electrical stimulation (Fig. 2(a)), all four RGC types are activated as long as they are close enough to the stimulation electrode. This results in an unnatural and even contradictory message communicated to the brain. In other words, due to the opposite situations that excite ON and OFF pathways (i.e., increment and decrement of light intensity, respectively) the signals sent to the brain following an electrical stimulation have a destructive effect on each other, resulting in creation of a vague or unstable perception in the brain [19], [20].

**Fig. 2:**
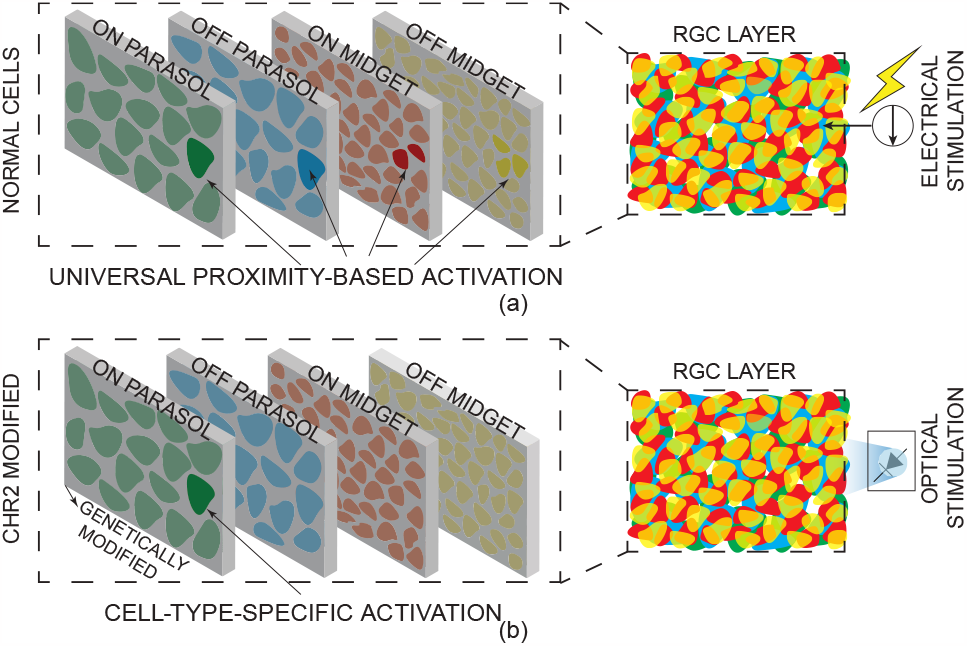
(a) Electrical stimulation evokes excitation in all RGC types in the proximity of the electrode. (b) Optical stimulation only evokes excitation in the genetically-modified cells and is capable of cell-type-specific excitation.

Selective activation of different RGCs using electrical stimulation is an open research topic. There are different approaches reported to bias the stimulation toward either ON or OFF pathways. Using smaller electrodes and pitch, manipulating the stimulus polarity, pulse duration, and frequency, and adding dedicated local return electrodes are some of the methods investigated for enhancing the selectivity of electrical stimulation [26]–[28]. Despite some level of success and perception improvements, these methods require calibration over time, making them less practical when considering the post-implantation variation [27] and usually achieved at expense of increased power consumption. Additionally, with electrical stimulation, charge balancing needs to be performed for each channel with a high precision considering the large number of electrodes, resulting in extra power consumption. On the other hand, through genetic modification, it is possible to make different retina cell types sensitive to distinct wavelengths. Therefore, as shown in Fig. 2(b), by generating different light wavelengths (e.g., different *μ*LED types embedded in an implantable array), selective activation of different RGC types can be achieved. It should be mentioned that currently, the specific promoter to genetically modify the ON bipolar cell types has been reported [7], [29]–[32], which is in line with our idea of placing the implant epi-retinal and stimulate the bipolar cells to take advantage of the remaining downstream retina network. However, a promoter to stimulate OFF bipolar cells has not been reported yet. Therefore, in this paper we have considered separate stimulation of ON and OFF bipolar cells, But also looked into the effect of stimulating only ON bipolar cells as that is the only viable possibility at the time.

## III. A Computational Model for Retinal Stimulation

To validate our hypothesis and to investigate the efficacy of optical stimulation as a potential treatment option for patients with retinal degeneration, we developed our model based on the ISETBio toolbox (Image System Engineering Toolbox-Biology) [33], which is a computational framework designed for calculating the properties of the front-end of biological visual systems and used in modeling the activation of retina layers for image formation. In doing so, the same biological properties of the retina from large-scale activity recording that has been used in ISETBio toolbox are considered with the exception that ON and OFF bipolar cells are assumed sensitive to different light wavelengths [33]. Fig. 3 shows different stages of retina cell layers’ activity for a given visual stimulus input and how each layer’s output instigates activity in the next layer. The toolbox itself is developed as an object-oriented model with different components of biological vision, each with its own biological and physical properties. Each of these objects are either defined as a MATLAB structure or a MATLAB class and the computations are implemented as transformations between these objects. As shown in Fig. 3(a), as the first step, the model extracts the radiance information of the input visual stimulus (defined as the “scene object”). Then, 31 points with a distance of 10nm are picked from the the visible light wavelength range from 400nm to 700nm, and the radiance information for each wavelength is calculated. Next, the spectral irradiance image at the cone photo-receptors (i.e., the “optical image object”) is computed by applying the physical properties of the eye (i.e., cornea, lens, pupil, and vitreous gel). The results show that, not only the number of photons that reach the photo-receptor layer are less than the visual stimulus, but the physical attributes of the eye also play a filtering role and attenuate the high-frequency content. As shown in Fig. 3(b), the photo-receptor layer is defined as “cone mosaic object”. The photo-receptor array’s absorption is calculated based on the spectral irradiance information of the optical image and the cone mosaic object’s biological and physical properties. This calculation is repeated several times (i.e., sequences), and each time a random eye movement is applied to account for the natural micro-saccade movements in a healthy eye. The “cone mosaic object” incorporates the three types of cone photo-receptors (i.e., L, M, and S type). It is possible to control the distribution and properties of each type, which allows for setting them in a way that matches the fovea (i.e., high photo-receptor-density region in retina) or periphery (i.e., low photo-receptor-density region). The generated photocurrent signals (proportional to the absorption levels shown in Fig. 3(b)) by the photo-receptors is used to calculate level of activation in “bipolar cells object” (shown in Fig. 3(c), left) for different types of bipolar cells (i.e., ON and OFF diffuse, and ON and OFF midgets). The average current (i.e., activity) in the bipolar layer shows that the ON and OFF cells activities peak at sections of the visual stimulus that are complementary to each other. As the final step, the bipolar cells’ output current is converted to the RGCs spiking pattern of the “RGC object”. Similar to the bipolar layer, for the RGC layer, four main types are RGC cells are taken into account (i.e., ON and OFF parasols, and ON and OFF midgets). Their activity is shown in Fig. 3(c) as the spiking density, which is the normalized number of generated spikes. The figure shows that, similar to the bipolar layer, ON and OFF RGCs have their peak of activity at complementary points of the visual stimulus. The RGC spiking pattern is generated based on a activation model, which can be a linear, a generalized linear (GLM), a linear-nonlinear Poisson, or a coupled GLM model [34].

**Fig. 3:**
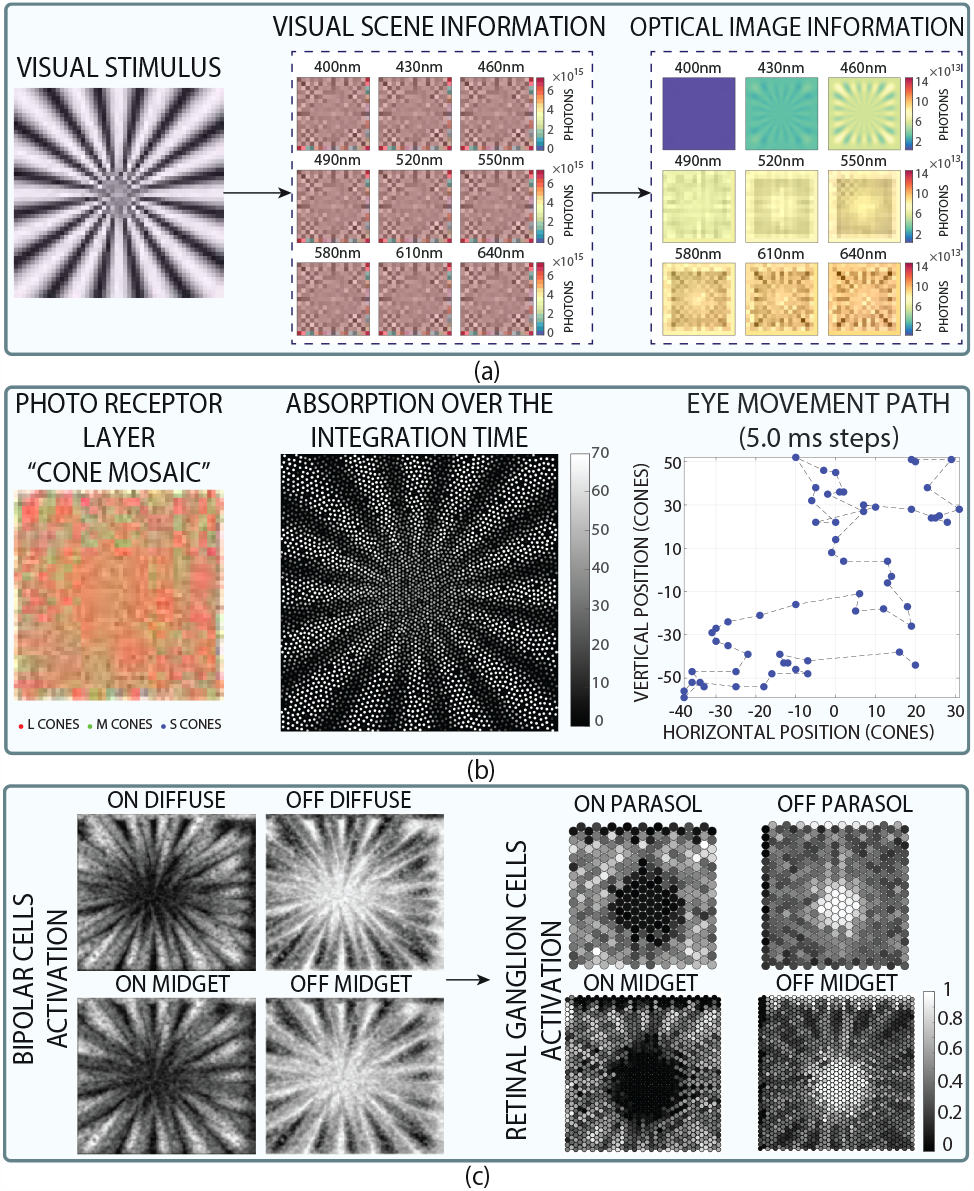
A step-by-step example showing how a visual stimulus input is fed to and processed by the developed ISETBio-based model at different stages from photo-receptor cells all the way to the spiking patterns generated in the RGCs to be sent to the brain.

In [19], the authors have developed an ISETBio-based model to generate the visual perception resulted from activation of bipolar cell layer in response to electrical stimulation (i.e., bypassing the photo-receptor layer). In their model, the electrical stimulation parameters (i.e., spatial and temporal intensity) are defined based on the visual stimulus captured by a camera. The patient’s perception estimation through linear reconstruction of retinal ganglion cells activation is also included in their model which can be used to study the adjustment of different stimulation parameters.

In this work, we have developed a an ISETBio-based framework that translates the captured visual stimulus into optical stimulation pattern that are used for activation of genetically-modified bipolar cells, with each pathway being transfected with a distinct opsin. By doing so, the ON and OFF pathways were made sensitive to visual stimulus irradiance increment and decrement, respectively. In this model in order to generates the temporal and spatial activation pattern of different retina cell layers (i,e,. bipolar cells), the input visual stimulus must be a sequence of images. Accordingly, because the activation triggers by either increment or decrement in the irradiance of the input visual stimulus, a sequence of constant images would not work well as the input. To solve this challenge, a random movement across both X and Y axes is introduced, to generate the sequence of images fed to the model. This random eye-movement mimics the natural micro-saccade movement in a healthy eye. After defining the stimulation pattern in the bipolar cells, the rest of the downstream network is assumed functional and used to calculate the spiking activities of different RGCs.The patient’s perception estimation is reconstructed using linear model based on the induced activities of RGCs. Beside the pathway-specific stimulation, this model also allows for defining the optical stimulation parameters (e.g., intensity, frequency, and duty cycle) and the *μ*LED array physical specification (e.g., *μ*LED array spatial resolution, spatial coverage, placement, and light divergence). This is used to study the effect of these adjustments on the reconstructed perception and to optimize the parameters while considering the area and power restrictions of an implantable neuro-stimulator device.

### A. Stimulus to Inference with Electrical Stimulation

To generate a baseline for comparison, we first regenerated the response (i.e., visual perception) to electrical stimulation based on what is developed in [19]. Fig. 4(a) shows the spiking activity density of different RGC types in response to electrical stimulation of bipolar cells assuming natural function for the rest of retina downstream network. As expected, due to the indiscriminate activation of electrical stimulation, the ON and OFF cells have similar spiking density, which is higher in the brighter areas of the visual stimulus and lower in the darker regions. This is contrary to what happens in a healthy retina, where these cells spiking activities peak at opposite light intensities of the input visual stimulus. Therefore, two contradicting messages are communicated by the ON and OFF pathways in response to the electrical stimulation, which combine destructively during linear reconstruction of the image. Consequently, the estimated perception is only marginally more correlated to the visual stimulus than noise, which is the same outcome also reported in [19] and [18].

**Fig. 4:**
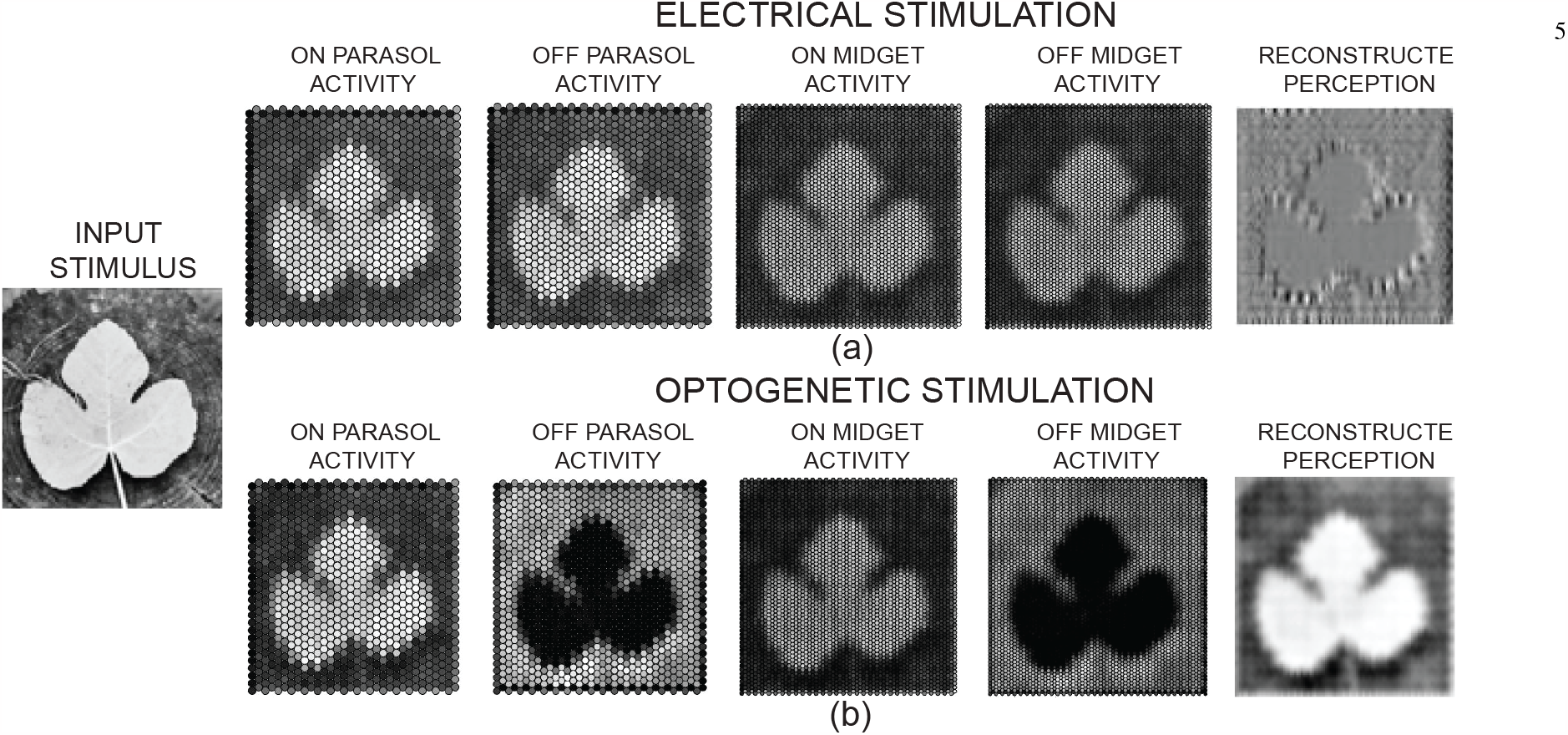
Simulation results showing the input stimulus (left), the four types of RGCs spiking activity density and their linear reconstruction for (a) electrical stimulation, and (b) optogenetic stimulation.

### B. Stimulus to Inference with Optical Stimulation

Fig. 4(b) shows the spiking activity density in RGCs in response to the same visual stimulus that is translated into optical stimulation pattern for bipolar cell layer and the natural downstrean network is assumed for defining the RGCs activities. The optical stimulation employs two distinct optical wavelengths for ON and OFF pathways, which respond to increment and decrement of light intensity. In practice, this can be done by employing a dual-color *μ*LED array, where the ON pathway cells are genetically modified by a blue-light-sensitive opsin (e.g. Chronos [35]) and the OFF pathway cells are genetically modified by a red-shifted opsin (e.g. Chrimson [35]). It should be noted that these two opsin light sensitivity wavelength lies within the visible range. However, considering that our current target is a fully degenerated retina, there is no constraint about the remaining healthy photoreceptors. Under these circumstances, the ON and OFF pathways cells exhibit distinctly different activations, same as what happens in a healthy retina. As a result, as shown in Fig. 4(b), the estimated perception calculated using linear reconstruction is highly resemblant to the visual stimulus.

To further compare the electrical and optical stimulations, we monitored the spiking activity of different RGCs in response to the two types of stimulation over time (presented in Fig. 5). The figure shows that for the electrical stimulation, all cells located in a close proximity of the stimulating electrode, regardless of their type, have a similar spiking activity. However, for the optogenetic stimulation, the cells from ON and OFF pathways have clearly different spiking activities. Considering the fact that both bipolar cell types (midget and diffuse) from each pathway are made sensitive to the same wavelength, the activation in the following RGCs are similar. They are only differentiated by the fact that the parasol cells have larger size and receptive fields than the midget cells, therefore, the plots show a denser spiking activity for midget cells due to their higher count in the same area, in comparison with the parasol cells.

**Fig. 5:**
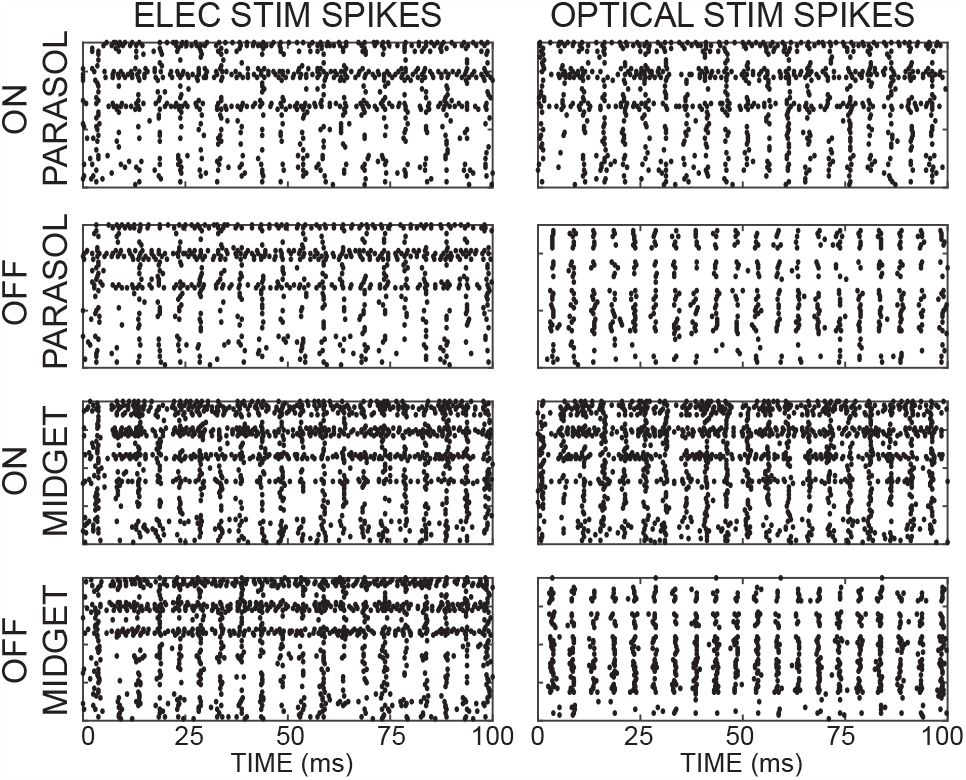
Simulation results showing the spiking activity of four RGC types over time, highlighting the distinction between ON and OFF pathways in optical stimulation, which is absent in electrical stimulation.

## IV. Analysis and Optimization of Stimulation parameters and *μ*LED Array Specifications

In this section, the effect of different optical stimulation parameters and the *μ*LED array’s physical specifications are investigated. We have also included a discussion on each simulation result and how they could be used for parameter optimization.

### A. μLED Array Spatial Resolution

The developed model is used to investigate the impact of increasing the optical stimulation’s *μ*LED array density in improving the visual perception, and also to compare that to the same *μ*electrode array density in electrical stimulation. This will compare the two modalities in terms of their potential for channel-count scalability as well. To do this, the same visual stimulus as in Fig. 4 is fed to the models for optical and electrical stimulation. Then the structural similarity of the reconstructed visual perception to the input visual stimulus is calculated and the results are normalized with respect to white noise’s (i.e., when random numbers are used for each pixel’s brightness with uniform distribution) structural similarity to the visual stimulus. The three terms that make up the structural similarity quality assessment index are brightness, contrast, and structure. The three terms are multiplied to create the overall index [36]. This experiment is repeated for 12 different *μ*-electrode/*μ*LED array densities. As shown in Fig 6 our simulation results made it evident that the perception degradation due to the cell-type-non-specific activation in electrical stimulation is so severe that negatively affect factors such as reducing electrode size/pitch. In contrast, the results indicate that increasing the *μ*LED array’s density yields a considerable improvement in the visual perception. As a more visual example, Fig. 7 shows the linear reconstruction results for optogenetic and electrical stimulations with *μ*LED/*μ*electrode array of 17.5*μ*m, 35*μ*m, and 70*μ*m pitch, respectively. These results confirm the efficacy of increasing optical stimulation array density in improving the resulted visual perception in optogenetic stimulation, while no notable improvement is observed for electrical stimulation.

**Fig. 6:**
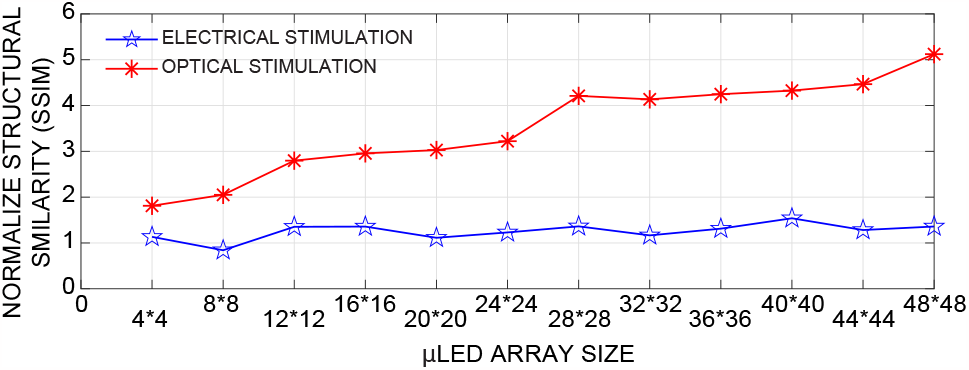
Simulation results comparing optical and electrical stimulation in terms of normalized structural similarity of the visual stimulus to the estimated perception as a function of the *μ*LED/*μ*electrode array density.

**Fig. 7:**
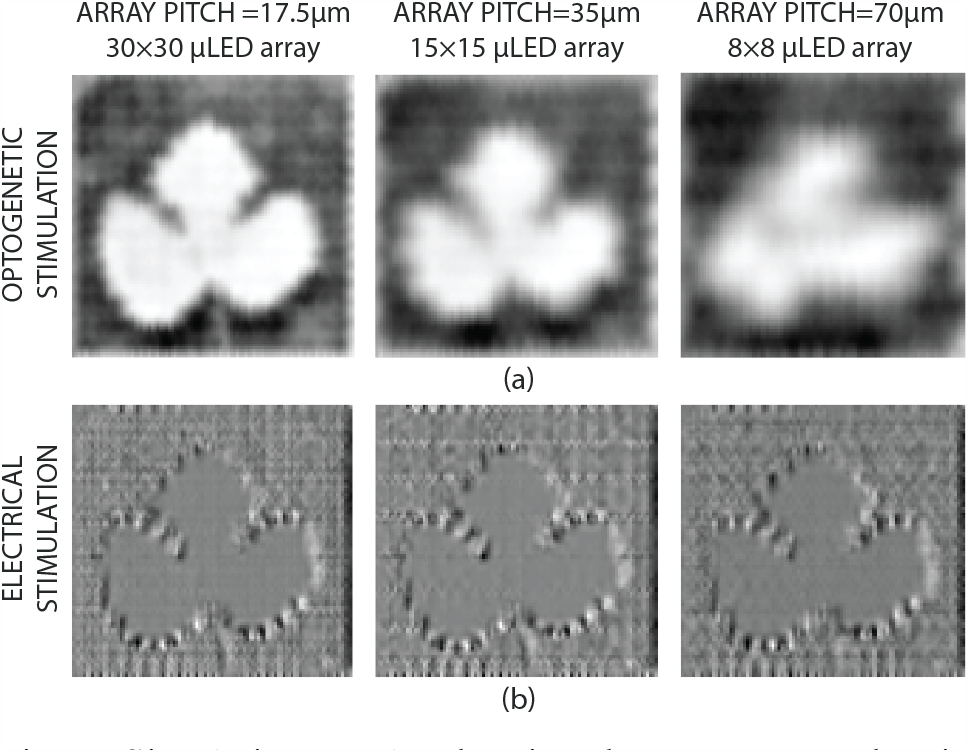
Simulation results showing the reconstructed estimated perception as a result of (a) optogenetic stimulation and (b) electrical stimulation for 17.5*μ*m, 35*μ*m, and 70*μ*m *μ*LED/*μ*electrode array pitch.

### B. μLED Array Spatial Placement

It is ideal for a stimulator designed to restore the functionality of a degenerated retina to be implanted where the density of bipolar cells (or RGCs, depending on the stimulation target) is at the highest. This is mainly to impact more neuronal pathways to the brain (i.e., higher spatial coverage), to maximize the stimulation efficacy. In the human eye’s retina, placing the stimulator’s *μ*electrode/*μ*LED array in a zone called fovea is ideal for this exact reason [37]. However, due to their indiscriminating activation, electrical stimulation electrodes are often placed peripheral to the fovea (e.g. in low-density areas such as raphe) to avoid unwanted cell stimulation [37]. This results in the stimulation to only affect parts of the retina that have a sparse cell population, further reducing the chance of success in effective vision restoration using this approach. Thanks to their cell-type specificity, this is not the case for optical stimulators and they can directly target areas with high cell-density.

We used the developed model to quantitatively illustrate the impact of placing an optical neuro-stimulator closer to the fovea, in achieving a better visual perception. Fig. 8 shows the normalized structural similarity of the reconstructed visual perception for a visual stimulus input with respect to white noise input at different distances from fovea. The figure clearly shows the importance of implant placement in achieving a better visual perception (same array size and light intensity used for all cases).

**Fig. 8:**
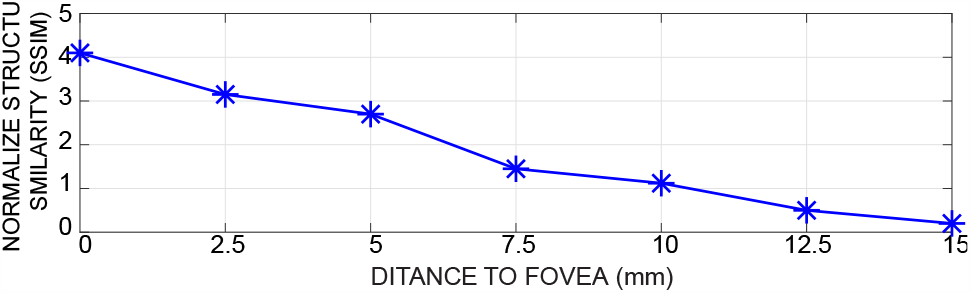
Simulation results showing the normalized structural similarity of the visual stimulus to the estimated perception as a function of distance to fovea.

This is more visually evident in the three example pictures shown in Fig. 9.

**Fig. 9:**
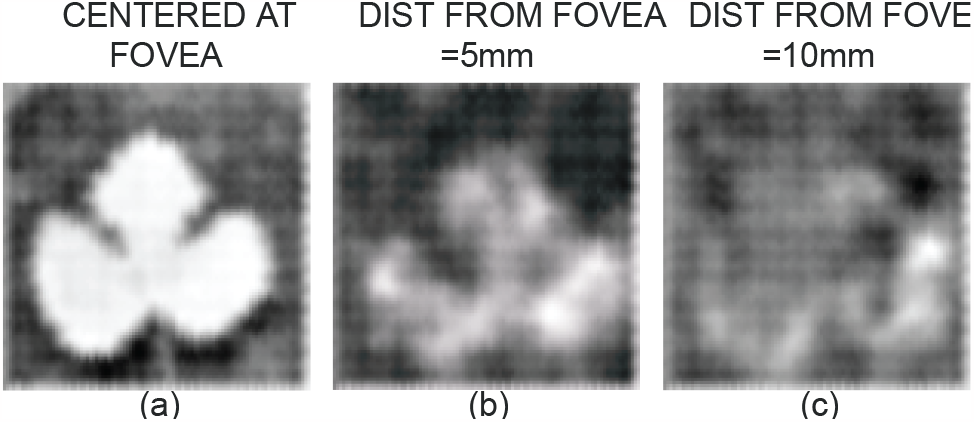
Simulation results showing the reconstructed estimated perception as a result of optical stimulation when the implant is (a) centered at fovea (b) at a 5mm distance from fovea, and (c) at a 10mm distance from fovea.

### C. μLED Array Light Divergence

A practical concern often cited about implantable *μ*LEDs used for optogenetic stimulation is their inherently-divergent light beams [38]. The main problem is that the poor light directivity leads to a poor spatial resolution for the optical stimulation, hence the LED light activates the nearby cells. While optogenetics ensures that only a specific type of cell is activated by the optical stimulation, it cannot stop the neighboring cells from being activated if they are of the same type as the target cells (i.e., similarly-transfected). Fortunately, in human eye’s retina, it has been shown that while the neighbor cells from different types have little to no correlation, the neighbor cells of the same type are highly correlated [34], [39], [40]. This is mainly due to the fact that these cells are usually affected by similar inputs from the previous retina layers. Therefore, the light divergence, at the microscopic level (i.e., multi-cell activation instead of a single-cell activation) can be even beneficial as it engages more relevant cells in the information transfer path.

The above discussion is not valid at the macroscopic level, meaning that cells of the same type that are far from each other are likely to be uncorrelated, hence, should not accidentally get activated together. This suggests that there is an optimal radiation field for the *μ*LEDs that results in the highest visual perception quality. Since the radiation field depends on both the *μ*LEDs’ spacing (i.e., the pitch) and their divergence angle, to investigate this optimal point, we first introduce a parameter called the divergence factor as,

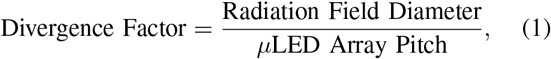

where the *μ*LED’s radiation field diameter is defined as the diameter of the surface area at the targeted distance from the optical source that all the transfected neurons inside it are stimulated by the optical illumination of that *μ*LED; and the *μ*LED array pitch is defined as the physical distance between the centers of the two neighboring *μ*LEDs in an array. In order to model the effect of light divergence, a spatial filter is applied to the optical stimulation pattern. Fig. 10 shows the spatial filters used to model different divergence factors, in which a Gaussian distribution is assumed for the *μ*LEDs radiation. For a divergence factor *<*1 (e.g., 0.5, as shown in Fig. 10), the radiation fields do not extend sufficiently to cover the entire distance between neighboring *μ*LEDs, which results in blind spots (i.e., not stimulated areas). On the other hand, for a divergence factor *>*1 (e.g., 2, as shown in Fig. 10), the radiation patterns from neighboring *μ*LEDs interfere, resulting in some of the cells being affected by more than one *μ*LED.

**Fig. 10:**
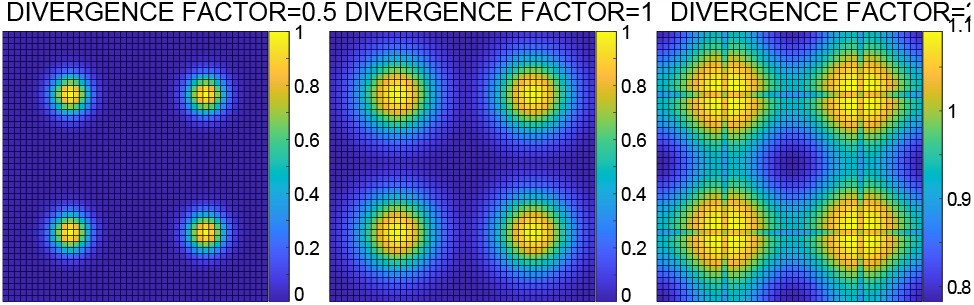
The spatial filters applied to the radiation of a 2 *×* 2 array of *μ*LEDs to implement divergence factors of 0.5, 1, and 2, assuming a 2-D Gaussian-distributed illumination, centered at each *μ*LED.

Fig. 11(a) shows the reconstructed visual perception for three different divergence factors, at two different *μ*LED array pitches. The figure confirms that the best result is for divergence factor of 1, while higher divergence factors result in blurring the perception (due to sending wrong messages to the brain by accidentally co-activating opsins that are far from each other), and lower values result in a fewer number of cells being activated, hence a proportional drop in the reconstructed perception’s quality. Fig. 11(b) shows the normalized structural similarity of a particular visual stimulus (same as the one in Fig. 5) with respect to a white-noise visual stimulus, at different divergence factors and for two different *μ*LED array pitches. The results confirm that regardless of the *μ*LED array pitch, the highest structural similarity is achieved for a unity divergence factor. These simulation results imply that for a specific *μ*LED that has a known radiation field, the optimum pitch for the *μ*LED array would be equal to the diameter of the radiation field. Fig. 12 shows a 3D plot that illustrates the importance of divergence factor for different distances between the implanted optical source and the fovea. As expected, for all distance values, a unity divergence factor yields an optimal performance. However, the figure suggests that divergence factor optimization becomes more important as the implant gets closer to the fovea. Our interpretation is that this is due to the higher density of neurons closer to the fovea, which results in a more significant negative impact of an interference between the *μ*LEDs’ radiation fields.

**Fig. 11:**
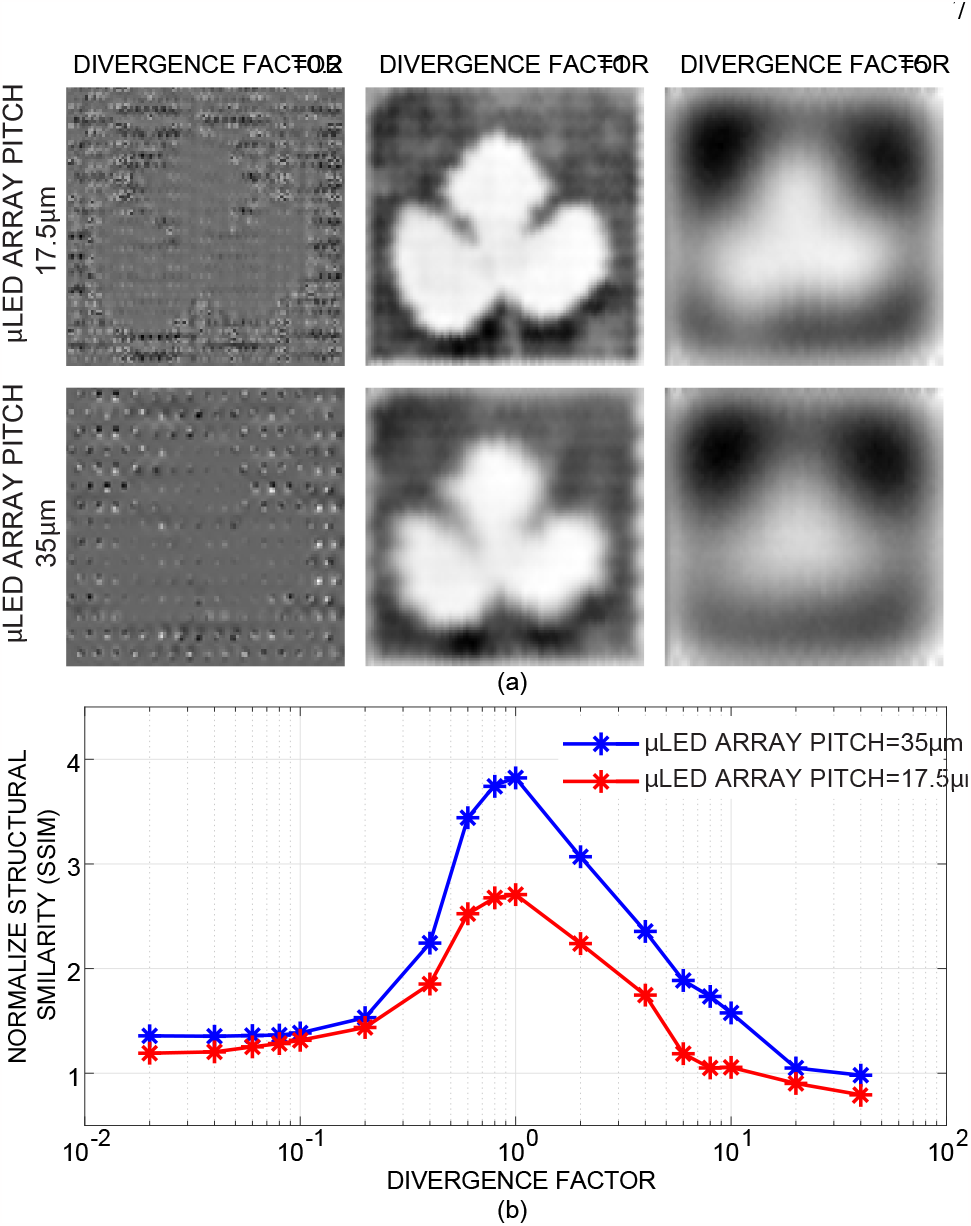
(a) Simulation results showing the reconstructed estimated perception for divergence factors of 0.2, 1, and 5 for *μ*LED array pitches of 17.5*μ*m and 35*μ*m. (b) Simulation results showing the normalized structural similarity of the visual stimulus to the estimated perception as a function of divergence factor for *μ*LED array pitches of 17.5

**Fig. 12:**
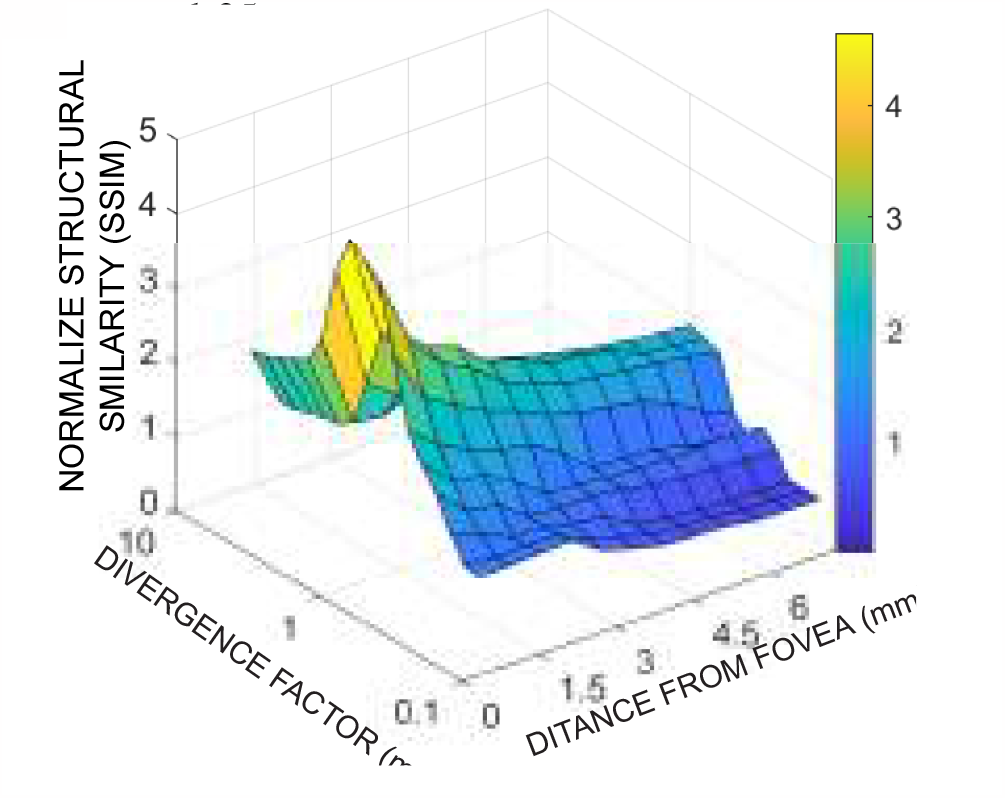
Simulation results showing the normalized structural similarity of the visual stimulus to the estimated perception as a function of divergence factor at different distances from fovea.

### D. Optical Stimulation Light Intensity

The instantaneous power required for driving the stimulating *μ*LEDs is orders of magnitude larger than all other blocks on an inductively-powered implantable retinal prosthesis (e.g., data receiver, power management, signal processing, etc.). Therefore, it is of critical importance for the stimulation light intensity to be optimized to avoid a power dissipation more than the minimum required (to maintain a high energy efficiency for the device) while ensuring that all targeted opsins receive sufficient power needed for activation.

We varied optical stimulation intensity over a range of more than two orders of magnitude and compared the resulted visual perception with the stimulus input. As shown in Fig. 13(a), the perception’s quality clearly peaks at an optimal intensity and diminishes for higher and lower values. While the quality degradation was expected for lower light intensities (due to inactivation of some of the opsins), we observed that a higher-than-optimal intensity could also result in diminishing the visual perception’s quality, in addition to making the device less energy efficient. Fig. 13(b) illustrate the effect of non-optimal and optimal intensities with three visual examples. Our interpretation is that for the lower intensities, the visual perception has a lower contrast due to some opsins not receiving sufficient optical power to get activated. For the higher intensities, we assume that the quality degradation is due to saturation of the genetically-modified cells because of receiving too much power [41].

**Fig. 13:**
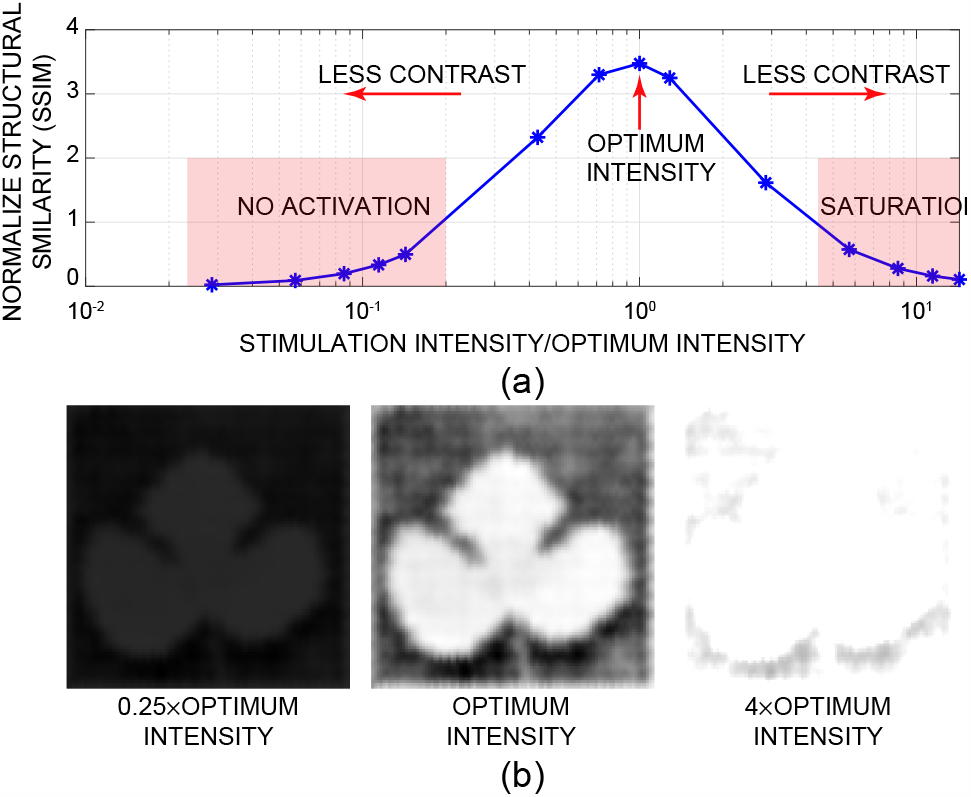
(a) Simulation results showing the normalized structural similarity of the visual stimulus to the estimated perception as a function of stimulation intensity divided by the optimum intensity. (b) Simulation results showing the reconstructed estimated perception for three different light intensities.

The above results indicate the importance of real-time neural activity recording for implantable optogenetic stimulators as also mentioned in [21]. Besides enabling closed-loop operation that could be used for diagnostic applications, real-time recording of neuronal activities (typically, electro-physiological recording) allows for calibrating the stimulation light intensity to avoid the above-mentioned problems at both sides of the spectrum.

### E. Single-wavelength Optical Stimulation

While implantable multi-wavelength (i.e., multi-color) *μ*LED arrays have been reported in literature [42], [43], custom micro-fabrication of a single-wavelength array poses far less complexity [42], [43]. Additionally, from an energy efficiency perspective, if we can conduct optical stimulation by turning on only one *μ*LED color, it could significantly (almost by 50%) reduce the required power consumption for stimulation.

To investigate this, we adjusted our developed model to only consider genetic modification for one of the pathways (i.e., ON or OFF). Fig. 14 shows the reconstructed perception due to optical stimulation of only ON or only OFF pathways. The results show that the above-mentioned fabrication and energy efficiency benefits come at the cost of quality degradation compared to a two-wavelength optical stimulation, but still a significantly better quality than the electrical stimulation. While in electrical stimulation the messages from two pathways had a destructive effect on each other, here only one pathway gets activated, hence, the message sent to the brain is still correlated with the stimulus input. Intuitively speaking, by using only one wavelength for optical stimulation, the optogenetic-based activation gain is reduced approximately by half, while the destructive effect of non-discriminate electrical stimulation is still avoided.

**Fig. 14:**
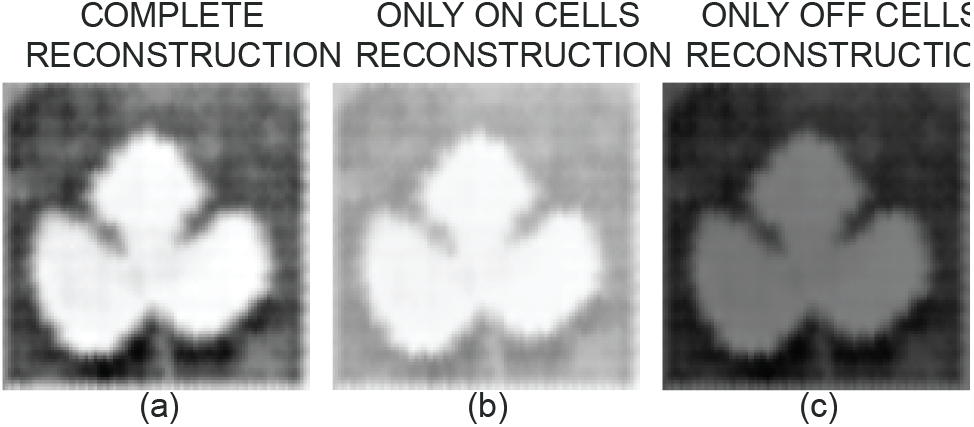
Simulation results showing the reconstructed estimated perception as a result of optical stimulation when (a) both ON and OFF pathways are activated, (b) only ON pathway is activated, and (c) only OFF pathway is activated.

Motivated by this hypothesis, we investigated if the lost activation gain in the absence of one of the pathways could be restored by increasing the stimulation intensity of the other pathway. Of course, this results in less energy benefit (or no energy benefit if we use 2*×* the intensity) for the single-wavelength case compared to a dual-wavelength stimulation, but it still has the advantage of less complex fabrication process. To do this, the normalized structural similarity of the reconstructed visual perception for the normal case (i.e., using both ON and OFF pathways) was compared with two activation cases of “only ON pathway” and “only OFF pathway”. For each of the three cases, we swept the stimulation light intensity by three orders of magnitude. As shown in Fig. 15 and as expected from the results of Section IV-D, there is an optimal intensity (i.e., *INT*_*OPT*_) for the normal case. The plots show that for the “only ON pathway” case, using *INT*_*OPT*_ results in 22% reduction in the perception quality, and increasing the intensity to 1.3*× INT*_*OPT*_ can only result in 2% improvement in the perception. An intensity beyond 1.3 *× INT*_*OPT*_ will have a negative impact on the reconstructed image quality. We assume this is due to opsins saturation. For the “only OFF pathway”, using *INT*_*OPT*_ will result in a more dramatic (i.e., 56%) quality loss, and the light intensity must be increased to 2.85 *× INT*_*OPT*_ to reduce the loss to 22%. We assume that this is mainly due to the higher activation threshold of the opsins used in the OFF pathway [44]. The above suggests that the simpler fabrication achieved by using one *μ*LED wavelength comes at the cost of at least 22% visual perception quality, which cannot be compensated even if higher power consumption than the “normal” case are tolerated.

**Fig. 15:**
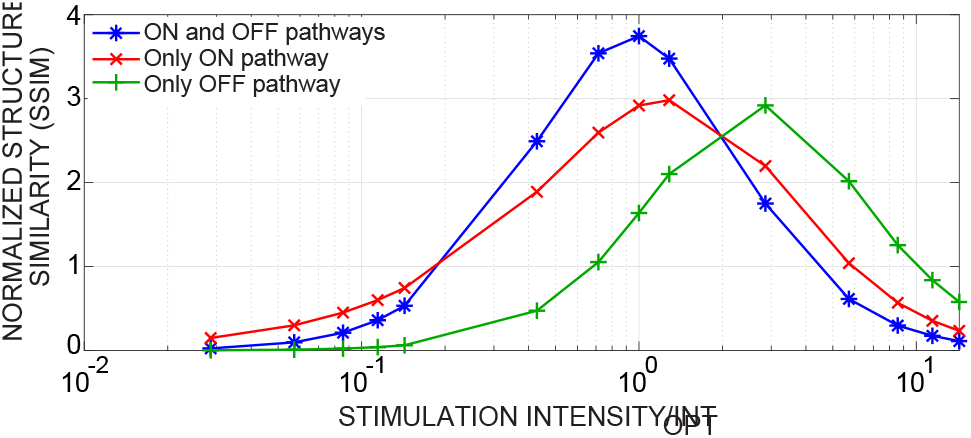
Simulation results showing the normalized structural similarity of the visual stimulus to the estimated perception as a function of stimulation intensity divided by the optimum intensity, when both ON and OFF pathways are activated (blue), only ON pathway is activated (red), and only OFF pathway is activated (green).

## V. Conclusion

We presented the design, validation, and simulation results of a retina-inspired computational framework that is developed to model the efficacy and optimize the implementation of an optogenetic epi-retinal neuro-stimulator. We discussed the potential of optogenetic stimulation as a treatment option for patients with retinal degeneration and presented fundamental advantages of an optical stimulator over an electrical one in terms of implantation invasiveness, system complexity, and most importantly, biological efficacy. Using the developed model, we showed the optical stimulation’s superiority, compared to electrical stimulation, in terms of the quality of the estimated visual perception it induces in the brain, and how this quality could be significantly improved by increasing the stimulator’s *μ*LED array resolution. We also used the model to investigate the impact of various optical stimulation parameters (such as optical light intensity and single-wavelength stimulation) and the physical specifications of the *μ*LED array (i.e., spatial resolution, location, and light divergence). For each case, we discussed the underlying biological rationale that explains simulation results and showed how these results could be leveraged towards implementing an implantable optical neuro-stimulator with an optimal energy efficiency and stimulation efficacy.

## References

[1] Age-Related Macular Degeneration: Facts & Figures [https://www.brightfocus.org/macular/article/age-related-macular-facts-figures]. Accessed 1 March 2022.

[2] W.L. Wong et al., “Global prevalence of age-related macular degeneration and disease burden projection for 2020 and 2040: a systematic review and meta-analysis,” in The Lancet Global Health, vol. 2, no. 2, pp. e106–e116, 2014.

[3] L. Yue et al., “Retinal stimulation strategies to restore vision: Fundamentals and systems,” in Progress in retinal and eye research, vol. 53, pp. 21–47, 2016.

[4] L. Theogarajan, “Strategies for restoring vision to the blind: current and emerging technologies,” in Neuroscience letters, vol. 519, no. 2, pp. 129–133, 2012.

[5] B. Roska and J.A. Sahel “Restoring vision,” in Nature, vol. 557, no. 7705, pp. 359–367, 2018.

[6] V. Wang, A.E. Kuriyan, “Optoelectronic Devices for Vision Restoration,” in Current Ophthalmology Reports, vol. 8, no. 2, pp. 69–7, 2020.

[7] M.E. McClements et al., “Optogenetic Gene Therapy for the Degenerate Retina: Recent Advances. Frontiers in Neuroscience,” in Frontiers in Neuroscience, vol. 14, pp. 1187–1203, 2020.

[8] M. S. Humayun et al., “Visual perception in a blind subject with a chronic microelectronic retinal prosthesis,” in Vision research, vol. 43, no. 24, pp. 2573–2581, 2003.

[9] S. Klauke et al., “Stimulation with a wireless intraocular epiretinal implant elicits visual percepts in blind humans,” in Investigative ophthalmology & visual science, vol. 52, no. 1, pp. 449–455, 2011.

[10] M. Monge et al., “A fully intraocular high-density self-calibrating epiretinal prosthesis,” in IEEE transactions on biomedical circuits and systems, vol. 7, no. 6, pp. 747–760, 2013.

[11] J. H. Park et al., “1225-channel neuromorphic retinal-prosthesis SoC with localized temperature-regulation,” in IEEE Transactions on Biomedical Circuits and Systems, vol. 14, no. 6, pp. 1230–1240, 2020.

[12] S.K. Kelly et al., “A hermetic wireless subretinal neurostimulator for vision prostheses,” in IEEE transactions on biomedical engineering, vol. 58, no. 11, pp. 3197–3205, 2011.

[13] K. Stingl et al., “Artificial vision with wirelessly powered subretinal electronic implant alpha-IMS,” in Proceedings of the Royal Society B: Biological Sciences, vol. 280, no. 1757, pp. 20130077, 2013.

[14] A. Asher et al., “Image processing for a high-resolution optoelectronic retinal prosthesis,” in IEEE transactions on Biomedical Engineering,, vol. 54, no. 6, pp. 993–1004, 2007.

[15] Z. Yu et al., “Toward the next generation of retinal neuroprosthesis: visual computation with spikes,” in Engineering,, vol. 6, no. 4, pp. 449–461, 2020.

[16] S. Nirenberg et al., “Retinal prosthetic strategy with the capacity to restore normal vision,” in Proceedings of the National Academy of Sciences, vol. 109, no. 37, pp. 15012–15017, 2012.

[17] N.L. Opie et al., ”Heating of the eye by a retinal prosthesis: modeling, cadaver and in vivo study,” in IEEE transactions on biomedical engineering, vol. 59, no. 2, pp. 339–345, 2011.

[18] B. Abbasi and J.F. Rizzo, “Advances in Neuroscience, Not Devices, Will Determine the Effectiveness of Visual Prostheses,” in Seminars in Ophthalmology, vol. 12, no. 3, pp. 1–8, 2021.

[19] J.R. Golden et al., “Simulation of visual perception and learning with a retinal prosthesis,” in journal of Neural Engineering, vol. 16, no. 2, pp. 025003, 2019.

[20] R. Iezzi et al., “Neurotransmitter-based retinal prosthesis modulation of retinal ganglion cell responses in-vivo,” in Investigative Ophthalmology and Visual Science, vol. 44, no. 13, pp. 5083–5083, 2003.

[21] T. Yousefi et al.,”An Energy-Efficient Optically-Enhanced Highly-Linear Implantable Wirelessly-Powered Bidirectional Optogenetic Neuro-Stimulator,” in IEEE Transactions on Biomedical Circuits and Systems, vol. 14, no. 6, pp. 1274–1286, 2020.

[22] D.D. Bezshlyakh et al., “Directly addressable GaN-based nano-LED arrays: fabrication and electro-optical characterization,” in Microsystems & Nanoengineering, vol. 6, no. 1, pp. 1–10, 2020.

[23] M. Asad et al., “Thermal and optical properties of high-density GaN micro-LED arrays on flexible substrates,” in Nano Energy, vol. 73, pp. 104724, 2020.

[24] T. Yousefi and H. Kassiri, “A Biologically-Informed Computational Framework for Pathway-Specific Spiking Patterns Generation and Efficacy Evaluation in Retinal Neurostimulators,” in IEEE Biomedical Circuits and Systems Conference (BioCAS), pp. 1–5, 2021.

[25] E. Richard et al., “Recognizing retinal ganglion cells in the dark,” in Advances in Neural Information Processing Systems, vol. 28, pp. 2476–2484, 2015.

[26] C. Sekirnjak et al., “High-resolution electrical stimulation of primate retina for epiretinal implant design,” in Journal of Neuroscience, vol. 28, no. 17, pp. 4446–4456, 2008.

[27] L.H. Jepson et al., “Focal electrical stimulation of major ganglion cell types in the primate retina for the design of visual prostheses,” in Journal of Neuroscience, vol. 33, no. 17, pp. 7194–7205, 2013.

[28] V.H. Fan et al., “Epiretinal stimulation with local returns enhances selectivity at cellular resolution,” in Journal of Neural Engineering, vol. 16, no. 2, pp. 025001, 2019.

[29] A. Berndt et al., “High-efficiency channelrhodopsins for fast neuronal stimulation at low light levels,” in Proceedings of the National Academy of Sciences, vol. 108, no. 18, pp. 7595–7600, 2020.

[30] Z. H. Pan et al., “Optogenetic approaches to restoring vision,” in Annual review of vision science, vol. 1, pp. 185–210, 2015.

[31] S. D. Klapper et al., “Biophysical properties of optogenetic tools and their application for vision restoration approaches,” in Frontiers in Systems Neuroscience, vol. 10, pp. 74, 2016.

[32] M. van Wyk et al., “Restoring the ON switch in blind retinas: Opto-mGluR6, a next-generation, cell-tailored optogenetic tool.,” in PLoS biology, vol. 13, no. 5, pp. e1002143, 2015.

[33] N.P. Cottaris et al., “A computational-observer model of spatial contrast sensitivity: Effects of wave-front-based optics, cone-mosaic structure, and inference engine,” in Journal of vision, vol. 19, no. 4, pp. 8–8, 2019.

[34] J.W. Pillow et al., “Spatio-temporal correlations and visual signaling in a complete neuronal population,” in Nature, vol. 454, no. 7207, pp. 995–999, 2008.

[35] N.C. Klapoetke et al., “Independent optical excitation of distinct neural populations,” in Nature methods, vol. 11, no. 3, pp. 338–346, 2014.

[36] Z. Wang et al., “Image quality assessment: from error visibility to structural similarity,” in IEEE transactions on image processing, vol. 13, no. 4, pp. 600–612, 2004.

[37] L. E. Grosberg et al., “Activation of ganglion cells and axon bundles using epiretinal electrical stimulation,” in Journal of neurophysiology, vol. 118, no. 3, pp. 1457–1471, 2017.

[38] T. Yousefi et al., “A 12.5 mg mm-Scale Inductively-Powered Light-Directivity-Enhanced Highly-Linear Bidirectional Optogenetic Neuro-Stimulator,” in IEEE Custom Integrated Circuits Conference (CICC), pp. 1–4, 2020.

[39] S.A. Baccus, “Timing and computation in inner retinal circuitry,” in Annu. Rev. Physiol., vol. 69, pp. 271–290, 2007.

[40] M. Vidne et al., “Modeling the impact of common noise inputs on the network activity of retinal ganglion cells,” in Journal of computational neuroscience, vol. 33, no. 1, pp. 97–121, 2012.

[41] N. Grossman et al., “Modeling study of the light stimulation of a neuron cell with channelrhodopsin-2 mutants,” in IEEE Transactions on Biomedical Engineering, vol. 58, no. 6, pp. 1742–1751, 2011.

[42] C. M. Kang et al., “Fabrication of a vertically-stacked passive-matrix micro-LED array structure for a dual color display,” in Optics Express, vol. 25, no. 3, pp. 2489–2495, 2017.

[43] S. Zhao et al., “Molecular beam epitaxy of III-nitride nanowires: Emerging applications from deep-ultraviolet light emitters and micro-LEDs to artificial photosynthesis,” in IEEE Nanotechnology Magazine, vol. 13, no. 2, pp. 6–16, 2019.

[44] F. Soto et al., “Efficient coding by midget and parasol ganglion cells in the human retina,” in Neuron, vol. 107, no. 4, pp. 656–666, 2020.

